# Biomechanical resilience of the femoral neck post-remission of Cushing’s Syndrome: a comparative analysis using QCT-based Finite Element models

**DOI:** 10.1101/2024.12.09.627485

**Authors:** Eduardo Soudah, Agustina Giuliodori, Joaquín A. Hernández

## Abstract

This study aims at evaluating femoral bone-related mechanical properties in female patients with long-term remission of Cushing’s Syndrome (CS). Sixty-four female subjects were included in this study and stratified in two groups: (a) 32 long-term remission of CS patients, and (b) 32 healthy (paired) control subjects. Quantitative Computed Tomography (QCT) was used to derive patient-specific Finite Element (FE) models. A sideways-fall impact was simulated for each subject and the resulting stress and strain values were compare among the two different groups. Our findings indicate that women with CS in remission exhibit impaired biomechanical properties in the femoral neck compared to controls, suggesting compromised bone properties in this population.

## 1. Introduction

Cushing’s syndrome is a rare endocrine disease characterized by cortisol hyper-secretion, induced mainly by a pituitary tumor or, less often, by an adrenal or ectopic neuroendocine tumor [1]. Regardless of the cause, chronic cortisol hyper-secretion can lead to a number of issues, including hypertension, central obesity and diabetes mellitus, as well as psychological dysfunction, hirsutism, muscle weakness and osteoporosis [2]. Since adequate diagnosis is often delayed 2–5 years, hypercortisolism exerts its harmful effect for a long time before it is diagnosed and treated [3]. Evidence from recent decades shows that, after effective treatment of CS, usually obtained by the surgical removal of the tumor, the normalization of cortisol secretion is not constantly followed by the complete remission of the associated clinical complications [1, 4]. It follows that, elucidation of the multiple pathogenetic mechanisms as well as long-term follow-up of these patients is essential in order to enhance the treatment of this complex endocrine disorder [5, 3].

One among the several clinical manifestations in patients who have been successfully treated is persistent bone disease [3, 4, 6, 7, 8]. During the active phase of CS, glucocorticoid excess exerts multiple and complex effects on the skeletal system, leading to bone loss and skeletal fragility [9, 10, 11, 12, 1]. Hypercortisolism interferes with both bone matrix formation and bone mineralization, either directly by impairing osteoblasts, osteocytes and osteoclasts, and, in an indirect way, by affecting parathyroid function and calcium, phosphorus, and vitamin D metabolisms [10, 1]. There exists evidence that bone loss induced by excess cortisol is most pronounced in skeletal sites with predominantly trabecular bone, such as the lumbar spine and femoral neck [13], which may be explained by the fact that bone remodeling occurs at bone surfaces, and trabecular bone has greater surface-to-volume ratio than cortical bone [14].

Active CS leads to non-traumatic fractures in up to 76% of patients [12, 15], even in patients presenting areal Bone Mineral Density (aBMD) with normal or slightly low values [11, 16]. This disparity suggests that glucocorticoids are especially detrimental to bone microstructure, affecting bone quality and strength [11, 16]. Long-term follow-up studies have shown a progressive improvement of bone mass and mineralization between 6 and 13 years after surgery, with a full recovery of aBMD in most of the patients [13, 17, 18, 19]. However, some reports indicate that estrogen-sufficient women in long-term remission of CS, may not achieve full recovery of aBMD and osteocalcin levels (a bone formation marker), even 11 years after remission compared to controls [6].

The absence of consensus on whether bone mass fully recovers after successful CS treatment, combined with the lack of long-term investigations into the evolution of bone micro-architecture in this disease, underscores the need for further research in this area. Consequently, our study aims to focus on evaluating the mechanical properties of the bone rather than analyzing densitometric measurements typically used to estimate bone loss. For this purpose, we employ Quantitative Computed Tomography (QCT), which offers a detailed volumetric assessment of trabecular and cortical bone, along with estimations of various biomechanical properties. Through the utilization of specialized calibration phantoms, the raw CT attenuation can be converted to volumetric Bone Mineral Density (vBMD), taking into consideration the bone mineral content within the analyzed bone volume [20, 21]. This density distribution enables the estimation of stiffness across the bone, with higher stiffness values attributed to regions with greater densities, such as cortical bone.

Due to the fact that QCT images enable the three-dimensional reconstruction and differentiation of anatomical bone structures, it is possible to build patient-specific QCT-based Finite Element (FE) models to evaluate the mechanical behavior of specific bone regions under particular boundary conditions [22, 23, 24, 25]. Depending on the assigned material and geometric properties, these models enable analysis of various scenarios, including bone deformations during falls, stress concentration in specific regions, or even bone fractures in severe cases (see, for instance, Refs. [26, 27, 28, 29]).

This study aims to evaluate bone-related mechanical properties at the proximal femur in female patients with long-term remission of CS, with a specific focus on analyzing strain and stress measures at the femoral neck. Patient-specific QCT-based FE models were generated for 64 female subjects, comprising 32 successfully treated CS patients and 32 healthy controls. Employing a numerical procedure simulating a sideways-fall impact, stress and strain distributions at the proximal femur were computed for each patient. Comparative analysis of stress/strain peak values at the femoral neck was conducted between CS patients in long-term remission and age-, Body Mass Index (BMI)-, and menopause-matched controls. The objective is to determine whether biomechanical abnormalities persist in CS treated patients despite remission.

## 2. Materials and methods

### 2.1 Study outline

For this study, 64 female participants were recruited and divided into two groups: (a) 32 patients with CS in remission for at least 3 years, and (b) 32 matched controls. Pairs were selected according to age, BMI, and menopausal status. All subjects were under the age of 65 to mitigate the potential impact of age-related mechanisms on bone properties. All participants gave written informed consent for the scientific use of the radiological explorations and associated clinical data. All study procedures were conducted in accordance with the guidelines approved by the Ethics Committee and the Declaration of Helsinki.

Each participant underwent QCT scans of the total hip. A sliced narrow neck (NN) analysis was conducted, where mechanical properties such as buckling ratio, cross-sectional area, and average cortical thickness were derived as the mean value across nine slices of the femoral neck. After performing the QCT scans, the image sequences were processed to delineate the boundaries of the proximal femur (segmentation). Once segmented, the 3D geometry of the femur was constructed and discretized into finite elements. The grey-level values from the CT images were then interpolated onto the elements of the FE mesh to define a patient-specific stiffness distribution. The simulation of a sideways fall impact allowed us to estimate strain and stress distributions on the proximal femur and peak values at the femoral neck. All the FE simulations were run using an in-house FE code.

Dependent t-tests for paired samples were then conducted in order to determine whether CS treated patients had similar mechanical properties than control subjects. Null hypothesis (H0) is that belonging to any group (CS remission or control subjects) have no effect over dependent variables. Rejection of H0 would indicate a relation between the dependent variable and the group, implying different behaviors. In all instances, statistical significance was defined as p-value< 0.05 (two-tailed test).

### 2.2. Image acquisition

QCT image sequences of the subjects’ total hip were acquired using a Phillips Brilliance 16 scanner (Mindways Software, Inc. Austin, TX, USA). The scanning acquisition parameters were chosen based on protocols routinely used in the clinical setting. A calibration phantom was included in the scanner for the appropriate densitometric conversion, allowing the determination of linear regression parameters (*α*_*ct*_ and *β*_*ct*_) for the conversion from X-Ray attenuation (*X*_*r*_), given in Hounsfield Units, to Computed Tomography (CT)-based density (*ρ*_*qct*_), measured in grams per cubic centimeter (g/cm3). This conversion was performed as follows [25]:

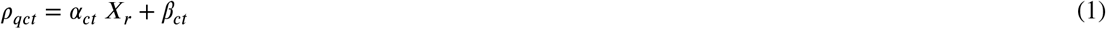

### 2.3. Patient-specific FE model

QCT image sequences acquired during the scanning process were imported into StradView [30] for the semi-automatic segmentation of the proximal femur region, combining methods like thresholding and manual selection. Subsequently, 2D contours were delineated for each image within the sequence, followed by the creation and exportation of a triangulated mesh representing the 3D surface of the femur [31]. This 3D surface was then imported into MATLAB (v. R2018a, The MathWorks, Inc.) for the automatic generation of the volumetric FE meshes, accomplished using a combination of meshing tools [32] and custom scripts.

The femoral bone was modeled as an elastic and heterogeneous material. The stiffness distribution was determined from densitometric measurements. To assign density values to each finite element in the mesh, an interpolation function, fitted based on CT image data, was used. Subsequently, a Young’s modulus value (E)[MPa] was assigned to each mesh element using a direct correlation between density and elasticity. This required the conversion of CT-based density (*ρ*_*qct*_) to physical density measures, namely ash (*ρ*_*ash*_) and apparent (*ρ*_*app*_) densities [21]. Based on a large review of literature given by Knowles et al. [21], we adopted the linear relationship *ρ*_*app*_ = *ρ*_*qct*_ and assumed a ratio between ash density and apparent density of *ρ*_*ash*_/*ρ*_*app*_ = 0.6 proposed by Dragomir-Daescu et al. [33]

Both cortical and trabecular bone tissues were modeled as linear isotropic elastic with Poisson’s ratio of 0.3. To calculate the local stiffness of the bone, we chose the following exponential law from the literature [33]:

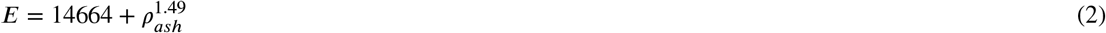

A tailored simulated sideways fall was conducted for each subject of the study, applying a distributed static load to the femoral head. The distal extremity of the femoral bone was fixed, while the external part of the greater trochanter was constrained in the direction of the force (see Fig. 1). The fall force resulting from a sideways fall was defined by the following equation:

**Figure 1:**
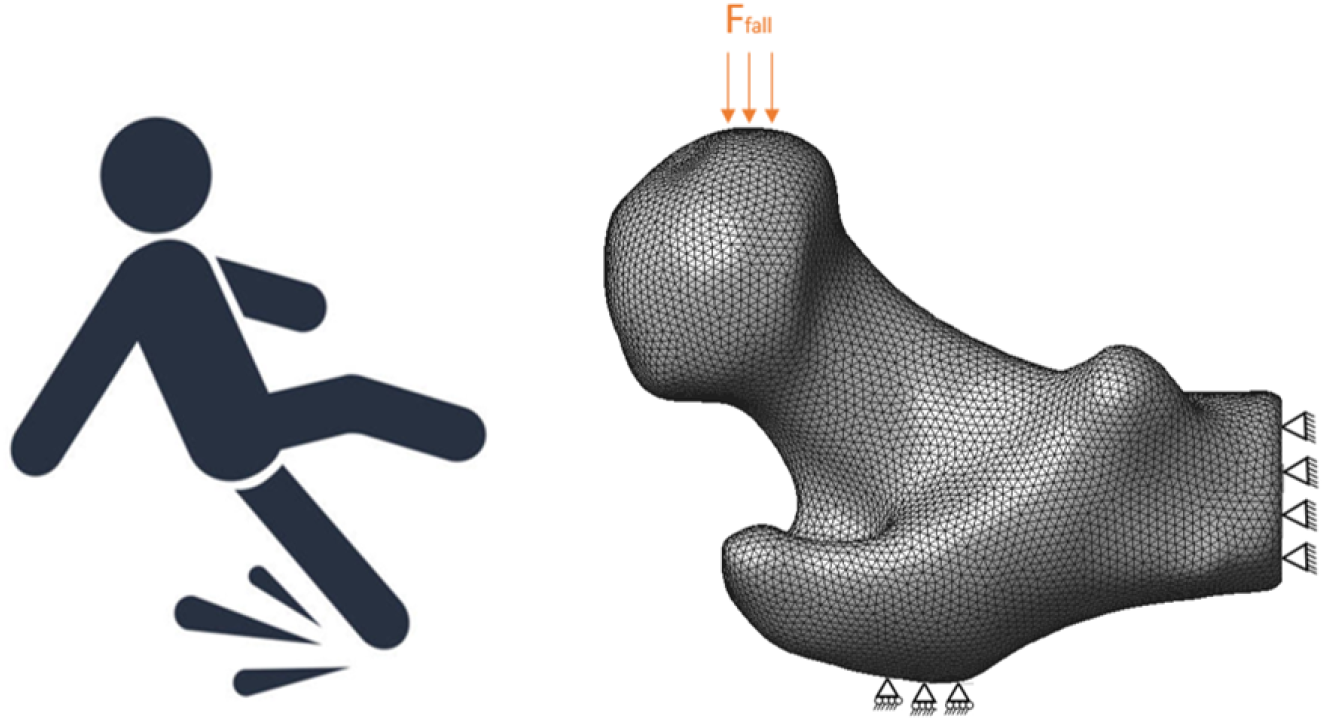
Boundary conditions for the FE simulation of patient-specific sideways-fall impacts.

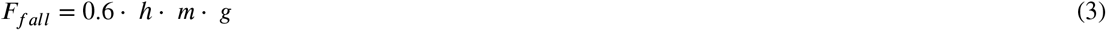

where *h* is the patient’s total height, *m* the patient’s mass and *g* = 9.81 *m*/*s*^2^ the gravity acceleration. The factor 0.6 is used to approximate the hip height relative to the patient’s total height, with the objective to evaluate the hip’s impact force during the sideways fall.

Four strain/stress parameters derived from FE simulation were calculated for each patient: Maximum Principal Strain (MPE), Maximum Principal Stress (MPS), Strain Energy Density (SED) and Von Mises stress (VM). Peak values were calculated at the femoral neck since it is the primary area for strain and stress concentration under this scenario (sideways fall). Mesh convergence analysis was carried out by incrementally increasing the number of elements to ensure the convergence of the results. The study determined that the voxel size is the most appropriate for accurate results. Thus, all FE meshes were generated taking into account the image’s voxel size.

## 3. Results

We begin by visually inspecting both stiffness and stress distributions on the proximal femur for each patient, as illustrated in Figure 2 for a selected subset. All results and data post-processing were obtained using the GID pre-post processing tool [32]. The images at the top display the (normalized) stiffness distribution, where higher stiffness in cortical bone regions is evident, as expected. At the bottom, the simulation results reveal how VM stresses mostly localize in the femoral neck for the given loading scenario. The patterns of stiffness and stress distribution do not appear to exhibit differences at first glance when comparing patient/control pairs.

**Figure 2:**
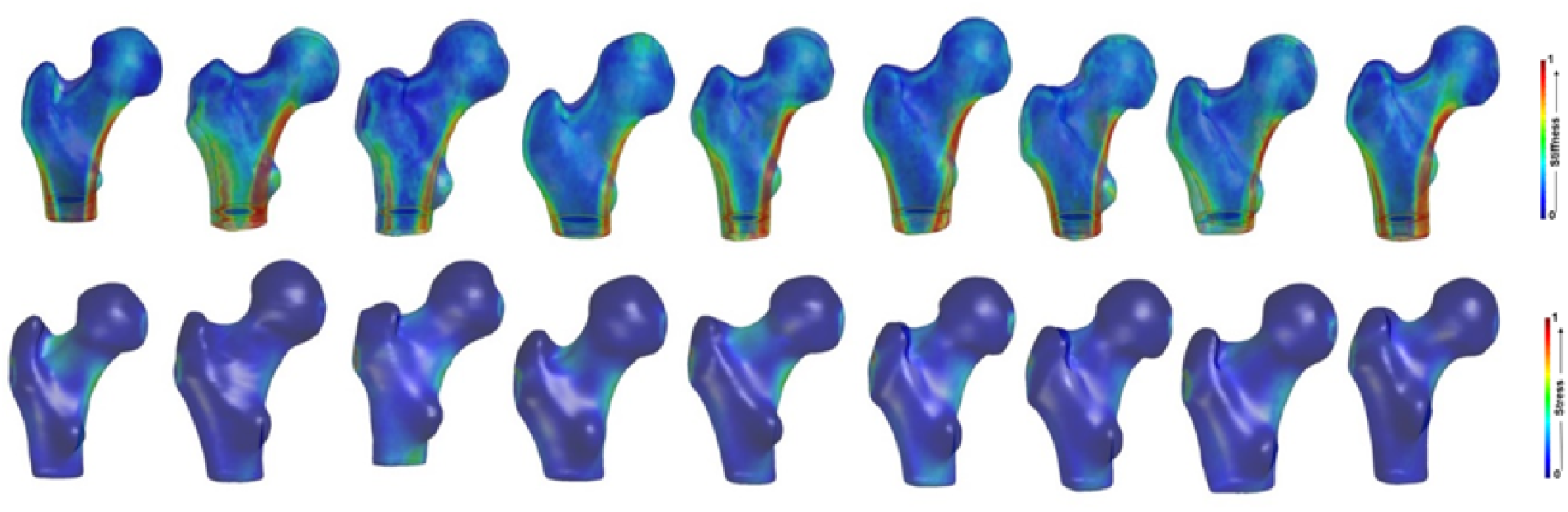
Normalized stiffness (above) and VM stress (bottom) distributions for a subset of patients.

Now we turn our attention to examine the mean stiffness values computed for both the proximal femur and femoral neck. In Figure 3, the box plots are divided by subject group, revealing that lower stiffness values are associated with treated patients compared to controls. Dependent t-tests indicate a statistically significant difference for the mean stiffness at the proximal femur (3588 ± 508 MPa for CS patients vs. 3877 ± 576 MPa for controls; *p* = 0 042), while the difference is not significant at the femoral neck (3701 ± 605 MPa for CS patients vs. 4027 ± 731 MPa for controls; *p* = 0.064).

**Figure 3:**
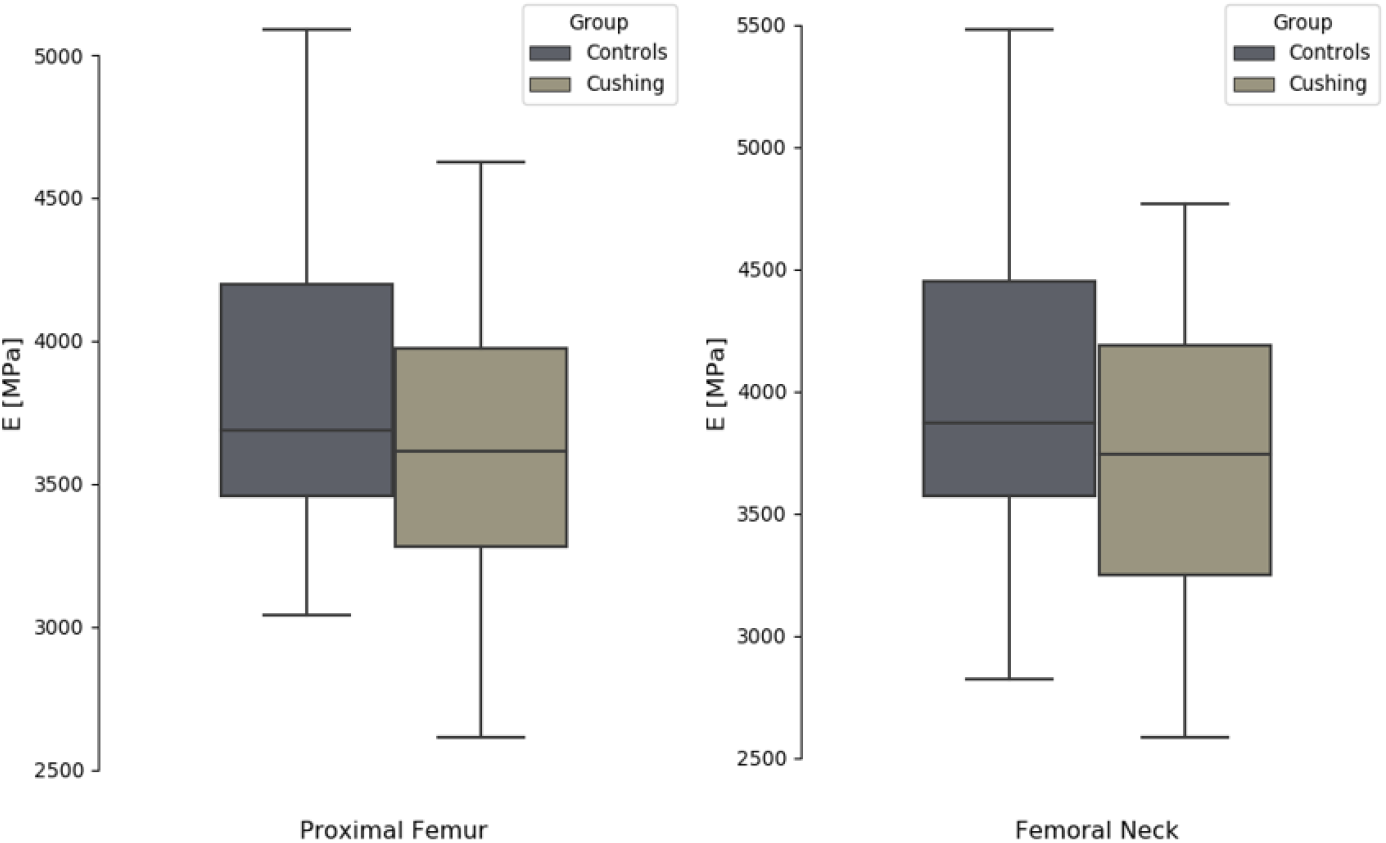
Box plots of mean stiffness values for proximal femur (left) and femoral neck (right) divided by groups.

The box plots included in Figure 4 illustrate the distributions per subject’s groups for biomechanical parameters derived from QCT. While cross-sectional area shows no significant differences between groups, both average cortical thickness and buckling ratio exhibit statistically significant disparities. Specifically, average cortical thickness is lower in treated patients compared to controls (0.24 ± 0.05 cm vs. 0.30 ± 0.08 cm, *p* = 0.001), whereas buckling ratio is higher in patients than controls (6.87 ± 1.98 vs. 5.5 ± 1.6, *p* = 0.005).

**Figure 4:**
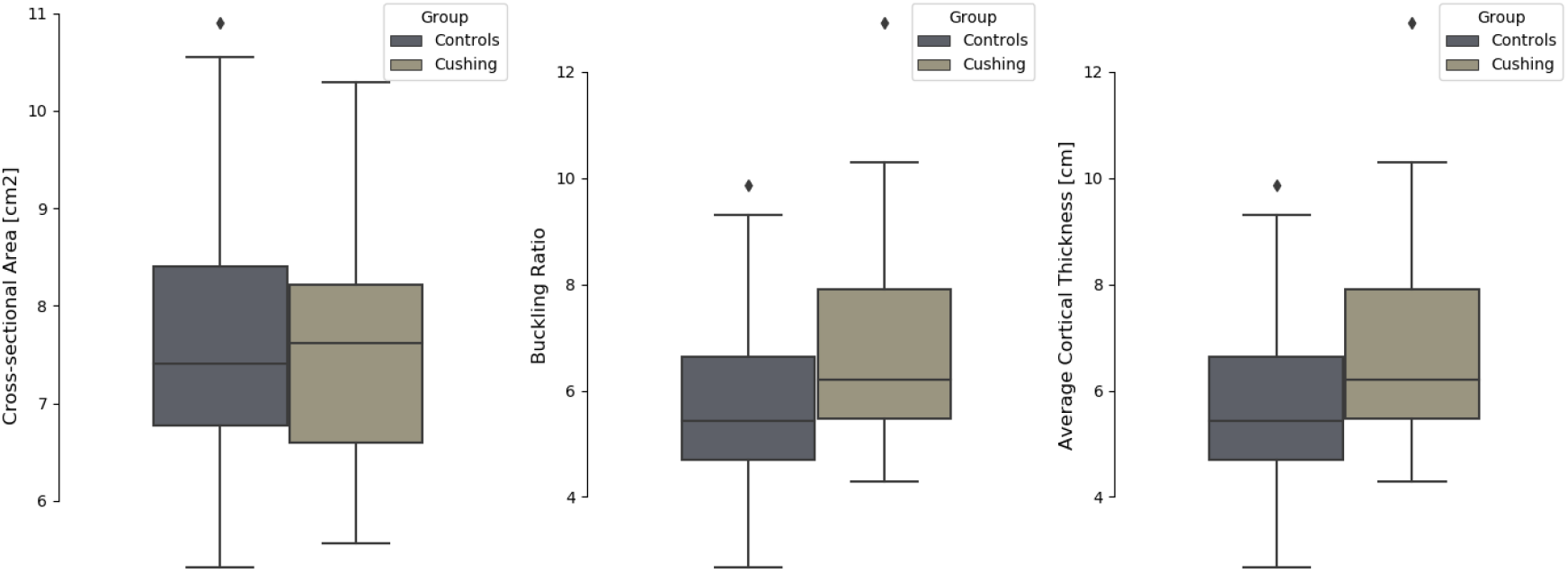
Box plots of biomechanical properties computed from QCT.

Figure 5 contains the box plots depicting the peak values of strain and stress measures at the femoral neck computed from FE sideways fall simulations. It is important to note that all strain and stress peak values exhibit significant differences between subject groups. Both MPE (0.035±0.007 for CS patients vs. 0.030±0.008 for controls; *p* = 0.015) and MPS (134.93 ± 36.38 MPa for CS patients vs. 114.99 ± 26.38 MPa for controls; *p* = 0.012), which reflect highest tensile/compressive normal strain and stress within the material, were elevated in treated patients compared to controls. The maximum SED, a measure of deformation energy per unit volume, was also greater in CS patients (0.92 ± 0.37 mJ/mm3 vs. 0.70 ± 0.25 mJ/mm3; *p* = 0.010). Finally, maximum values of VM stresses, measuring distortion energy, were higher in treated patients (126.18 ± 32.14 MPa vs. 108.12 ± 24.78 MPa; *p* = 0.010), suggesting increased susceptibility to failure for this group given a sideways fall scenario.

**Figure 5:**
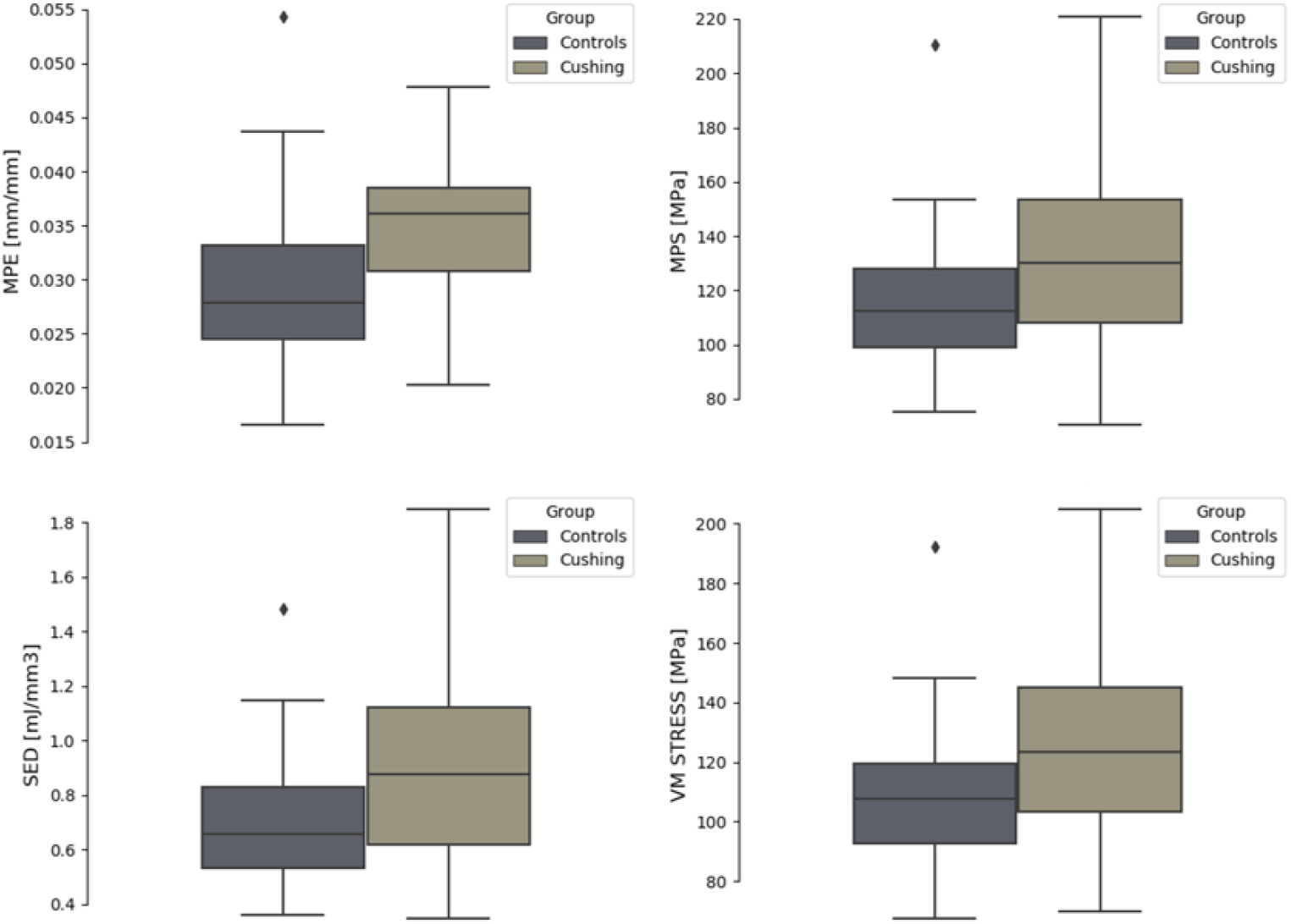
Box plots of peak strain/stress measures at the femoral neck divided by groups.

## 4. Discussion

The proximal femur of sixty-four female subjects were analyzed to assess whether bone mechanical properties of CS treated patients exhibits similar characteristics to that of control subjects. Stiffness distributions along with QCT-based biomechanical properties (average cortical thickness, buckling ratio, cross-sectional area) allowed for an initial comparison between subject groups. Simulating a sideways fall for each patient, whether in long-term CS remission or in the control group, enabled us to identify additional descriptors (MPE, MPS, SED and VM) that characterize the mechanical behavior of the femoral bone under such circumstances.

It is important to mention that in our FE models, we do not consider any elastoplastic/damage effects into the material behavior. Even we understood such considerations are crucial for understanding bone mechanical failure mechanisms, in this study we focus on how the distribution of cortical and trabecular bone influence on the stresses and strains during a sideways fall in patients with Cushing’s Syndrome. Moreover, although the intricacies of the trabecular bone geometry were not explicitly modeled, the densitometric measurements obtained from QCT were translated into corresponding stiffness values, providing a patient-specific distribution of the mechanical properties of the trabecular and cortical bone.

While stiffness distributions appear visually similar between the two groups, the mean stiffness values at the proximal femur are significantly lower for treated patients compared to controls. This suggests that bone mineral content does not fully recover after CS treatment, indicating persistent skeletal fragility. Given the lack of consensus on whether bone loss persists after long-term remission of CS, we believe that future investigations should leverage more advanced technologies such as High-Resolution QCT to assess bone micro-architecture in greater detail. This approach would be crucial for fully elucidating the underlying mechanisms that continue to affect bone mechanics following CS.

Our study has shown that, in terms of biomechanical properties, patients who received CS treatment has thinner average cortical thickness and, at the same time, higher values of buckling ratio. Note that, reduced cortical bone thickness is recognized to induce significant strains and stresses in femoral shafts, increasing the risk of hip fracture in elderly people [34]. In addition, buckling ratio is a measure of instability that captures the compensation mechanism of the femur by which it redistributes bone mass to counteract the bone loss in order to preserve bending strength [35]. The disparity in these parameters suggests that certain bone characteristics, which are not currently evaluated with standard clinical tools, may persistently be compromised in CS patients despite cortisol normalization.

Regarding FE-derived parameters, it is noteworthy that all strain and stress peak values at the femoral neck were significantly higher in CS patients compared to controls. It is important to acknowledge that in linear FE analysis, failure criteria are typically derived from the strain distribution since stresses vary with material properties, and plastic or damage variables are not considered in linear models [36]. Although our study does not directly assess failure, our findings suggest that strains at the femoral neck of treated patients may approach the threshold of mechanical bone failure earlier than controls under similar loading conditions. Other studies investigating bone mechanics in patients with osteoarthritis have found that, alongside cortical thinning, the loss of trabecular bone mass and connectivity contribute to skeletal fragility [37]. Thus, we believe that the elevated strain/stress values found for treated patients in our study may be associated with both cortical thinning and deterioration of bone material properties.

Finally, it is worth noting that our study did not explore age-related deterioration mechanisms that could also impact stiffness, and we are aware of the synergistic effect between increasing age and chronic GC therapy on bone mechanical properties [38, 39, 40]. However, our participants were paired according to age, and individuals older than 65 years were excluded from the study. Investigating whether older patients with CS remission are more susceptible to femoral fractures compared to age-matched controls who did not experience cortisol excess would be a valuable direction for future investigation.

## 5. Conclusions

This study demonstrates that QCT-based FE models of the proximal femur significantly improve the assessment of bone quality in CS patients by incorporating comprehensive descriptors of mechanical behavior. These descriptors consider patient specific geometric changes in cortical and trabecular bone, as well as the specific boundary conditions of the sideways fall. This more accurate bone assessment can help guide treatment decisions and improve patient outcomes in CS patients. Monitoring bone mechanics in these patients, even after long-term remission, remains crucial, and further research into the mechanisms influencing bone quality is essential.

## Acknowledgments

This work has received support from the Spanish Ministry of Economy and Competitiveness, through the ‘Severo Ochoa Programme for Centres of Excellence in R&D’ (CEX2018-000797-S). The authors also acknowledge the support of ‘MCIN/AEI/10.13039/501100011033/’ y por ‘FEDER una manera de hacer Europa’ (PID 2021-122518OB-I00).

